# Differential effects of piroxicam and nitroglycerine on memory and hippocampal neurochemistry in di-oestrous female rats

**DOI:** 10.64898/2026.06.30.735514

**Authors:** Folahan Joseph Kilanko, Bernard Omokheshi Adele

## Abstract

**Objectives:** To evaluate and compare the neuro-behavioural safety profiles of piroxicam and nitroglycerine by investigating their differential effects on cognitive function, spatial and recognition memory, and hippocampal neurochemistry in a di-oestrous female Wistar rat model.

**Methods:** Female Wistar rats at di-oestrous were randomly assigned to receive distilled water, piroxicam, or nitroglycerine orally for four consecutive days. Following treatment, spatial and recognition memory were evaluated using standard behavioural paradigms. Hippocampal tissues were analysed for acetylcholinesterase and glutamate activity, oxidative stress markers, and neuroinflammatory indices.

**Results:** Piroxicam improved recognition memory and was associated with increased glutamatergic activity and a compensatory rise in superoxide dismutase. However, it also elicited elevated nitric oxide signaling, lipid peroxidation, and localized neuroinflammatory markers in the hippocampus. In contrast, nitroglycerine impaired non-spatial memory during di-oestrous. Although both treatments preserved working memory, they produced distinct effects on object recognition, memory discrimination, oxidative stress parameters, and neuroinflammatory mediators.

**Conclusions:** Piroxicam and nitroglycerine exert differential effects on cognition and hippocampal neurochemistry during di-oestrous. Piroxicam improved recognition memory and produced distinct hippocampal neurochemical alterations, whereas nitroglycerine impaired recognition memory. These findings highlight the influence of menstrual pain therapeutics on cognitive function and hippocampal physiology under hormonally sensitive conditions.

## Introduction

Dysmenorrhea, a real challenge facing women of reproductive age regardless of race, age or socioeconomic background, affects 45-90 % of teenage girls and women [1]. It is typically characterised by painful menstruation, cramping pain in the lower abdomen that starts 1-2 days during menstruation and lasts for 2-3 days [2, 3]. Of the two forms of dysmenorrhea, primary dysmenorrhea is one of the most serious gynaecological conditions that affect young women [4]. Studies have shown that primary dysmenorrhea may be due to increased or abnormal uterine activity caused by uterine prostaglandins (PGs) produced by degenerating cells during endometrial sloughing [5]. Prostanglandins cause uterine myometrial hypercontractility and vasoconstriction [6]. People with primary dysmenorrhea have higher levels of PGF2α in their menstrual fluid correlating with the pain intensity [7].

Dysmenorrheic condition has been shown to lower quality of life and increase the risk of anxiety and depression [8]. According to Mesele et al. [9], occurrence of dysmenorrhea had a direct relationship with poor academic performance in female subjects. It was advanced that the accompanying pain caused difficulties in studying and sleep disturbances that negatively affect academic outcomes [10]. Memory function, the ability to retain and recall information [11, 12] may have disrupted in the dysmenorrheic subjects to result in the poor academic outcome. The role of PGs in memory deficit has been found positive in contextual fear conditioning while inhibition of cyclooxygenase (COX) activity disrupted contextual memory, thereby showing dual effect of PG in memory [13]. According to Hein et al. [13], both excessive and insufficient hippocampal prostaglandin levels have been reported to impair memory function as excessive hippocampal PGs level impaired working, spatial and long-term memory while inhibition of the cyclooxygenase pathway in the hippocampus disrupted contextual memory indicating that memory works best at moderate PG level. Nitroglycerine a nitric oxide donor, and piroxicam a common Nonsteroidal anti-inflammatory drug (NSAIDs), classified as prostaglandin synthetase inhibitors, are the commonly prescribed drugs to manage dysmenorrhea [14, 15]. This class of drugs work by inhibiting COX activity consequently causing reduction in prostaglandins level and associated cramps during menstruation. Since PG surge occurs during menstruation contributing to the dysmenorrheic experience, the use of COX inhibitors to manage dysmenorrhea may impact PG-modulated memory function. This study therefore investigated memory function in rats treated with piroxicam and nitroglycerine at di-oestrous.

## Materials and methods

### Experimental animals

#### Animals

Nulliparous female Wistar rats (weighing 120-140 g) were obtained at 6 weeks from the Animal House, Institute for Advanced Medical Research and Training, College of Medicine, University of Ibadan. The rats were acclimatised to standard laboratory conditions (22 ± 1 °C, 55 ± 5 % humidity), free access to normal mouse feed and filtered water, and a 12 h light/dark cycle (lights on at 07:00), for 14 days prior to the commencement of all experimental procedures. All experimental protocols followed the University of Ibadan’s Research Ethics Committee’s Animal Care and Use as well as the National Research Committee, 1996 for the Care and Use of Laboratory Animals, published by the National Academy Press, 2101 Constitution Ave. NW, Washington, DC 20055, USA. This study was approved by the University of Ibadan’s Research Ethics Committee’s Animal Care and Use (NHREC/UIACUREC/05/12/2022A).

### Model, grouping and drug treatment

The di-oestrous phase of female Wistar rats overlap with key pathophysiological aspects of primary dysmenorrhea. It is characterised by low oestrogen and progesterone, a hormonal environment associated with heightened inflammatory tone, increased prostaglandin sensitivity, and altered pain perception. This makes the di-oestrous phase a suitable biological window for modelling dysmenorrhoea-associated central effects.

A non-invasive and reliable method, vaginal cytology [16] was used to monitor and establish the oestrous cycle stages in the Wistar rats for 12 days, ensuring that the animals exhibited normal cycling patterns prior to the study. The presence, absence, or proportionate amounts of leucocytes, cornified (keratinized) cells, and epithelial cells (two types) defined the several oestrous stages (pro-oestrous, oestrous, met-oestrous, and di-oestrous). Animals in the di-oestrous phase were identified by the presence of predominant leukocytes in the vaginal smear [17].

Fifteen female Wistar rats, confirmed to be at their di-oestrous phase, were randomly grouped into three: Control (given distilled water, 0.3 mL) orally, Nitroglycerine (given 1 mg/kg nitroglycerine) orally and Piroxicam (given 3 mg/kg piroxicam) orally [18] for four days. The piroxicam, purchased under the brand name Feldene and Nitroglycerine were both obtained from Pfizer Pharmaceuticals, New York, NY, USA.

### Neuro-behavioural evaluation

#### Y-maze test

The Y-maze evaluated immediate spatial working memory based on the rats’ innate tendency to explore novel environments via spontaneous alternation behavior. Rats were acclimatised in the testing room for 1 min before testing. The maze was cleaned with 70 % ethanol between trials. EthoVision software recorded the animals for 8 min during arena exploration. An arm entry was counted when the rat’s entire body (excluding the tail) entered an arm. The spontaneous alternation percentage was calculated from the software reports to estimate short-term spatial working memory.

### Novel object recognition test (NORT)

NORT assessed non-spatial memory by exploiting the rats’ natural preference for novel objects over familiar ones. Rats were habituated to the empty arena for 5–10 min daily over 1–3 days to minimize handling stress.

During the training phase, two identical objects were placed at equal distances, and rats were allowed to explore them for 5–10 min. The rats were then returned to their home cages for specific delay intervals (5–10 min for short-term memory; 24 h for long-term memory). During the testing phase, one familiar object was replaced with a novel object, and rats explored the arena for 3–5 min. The time spent interacting with each object was tracked to calculate recognition and discrimination metrics:

Recognition Index (RI) was recorded as: Time spent on novel object -Time spent on familiar object

Discrimination Index (DI) was recorded as: (Time spent on novel object - Time spent on familiar object) / Total exploration time.

### Tissue collection

The rats were anaesthetised with ketamine and cervically dislocated. Thereafter, the skull was opened up to excise the whole brain which was rinsed in ice-cold phosphate buffered saline, blotted, weighed and quickly transferred over ice. Hippocampus was isolated over ice, weighed and homogenized in ice-cold 0.1 M phosphate buffered saline (pH 7.4). Supernatant collected over centrifugation (10,000 rpm, 4 °C, 10 min) was subjected to biochemical assays.

### Measurement of neurotransmitters, inflammatory cytokines and oxidative stress

Glutamate and acetylcholinesterase (AChE) levels were quantified in hippocampal tissue homogenate supernatants using commercially available enzymatic colorimetric assay kits (Nanjing Jiancheng Bioengineering Institute, Nanjing, China), following the manufacturers’ instructions. Briefly, samples were incubated with the appropriate reaction reagents at 37 °C, followed by substrate addition, and absorbance was measured using a microplate reader (SpectraMax M2, Molecular Devices, San Jose, CA, USA) at the specified wavelengths indicated by each assay protocol.

Levels of the pro-inflammatory markers, tumour necrosis factor-alpha (TNF-α) and myeloperoxidase (MPO) in hippocampal homogenate supernatants were determined using enzyme-linked immunosorbent assay (ELISA) kits (Neobioscience, Shenzhen, China). Nitric oxide (NO) production was assessed indirectly by measuring its stable metabolites (nitrite/nitrate) using a colorimetric assay based on the Griess reaction, according to the manufacturer’s instructions (Sigma-Aldrich, St. Louis, MO, USA).

Oxidative stress parameters, including malondialdehyde (MDA), superoxide dismutase (SOD), reduced glutathione (GSH), and hydrogen peroxide (H₂O₂), were quantified in hippocampal tissue homogenates using colorimetric assay kits (Nanjing Jiancheng Bioengineering Institute, Nanjing, China). All assays were performed strictly in accordance with the respective manufacturer’s protocols.

### Statistical analysis

Data were analysed using one way analysis of variance (ANOVA) and Tukey Post-hoc test for pairwise comparisons between groups at *P≤0.05* (Graphpad Prism Statistical Software, San Diego, CA, USA). Values were expressed as Mean ± SEM.

## Results

### Body and brain weight

Nitroglycerine and piroxicam treatment did not cause any significant change in the body weight, absolute brain weight, and relative brain weight when compared with control (Table 1).

**Table 1.** Body weight change, absolute and relative brain weight in control and treated rats.

| Parameters | Control | Nitroglycerine | Piroxicam |
| --- | --- | --- | --- |
| Body weight, g | 156.80 ± 8.32 | 139.80 ± 9.89 | 146.30 ± 5.54 |
| Absolute brain weight, g | 1.73 ± 0.05 | 1.65 ± 0.05 | 1.64 ± 0.01 |
| Relative brain weight, g | 1.11 ± 0.03 | 1.19 ± 0.07 | 1.12 ± 0.04 |

Note: Data are presented as Mean ± SEM (n = 5). Statistical significance is defined at $p \leq 0.05$ ; $P > 0.05$ compared with the control group.

### Influences of nitroglycerine and piroxicam on neuro-behavioural studies in di-oestrous rats

#### Y-maze test

Spontaneous Alternation in Nitroglycerine- and piroxicam-treated (65.55 ± 2.25 and 75.81 ± 4.33 %, respectively) did not differ significantly when compared with control (64.82 ± 1.48 %). It however increased in piroxicam-treated by 16.95 % and 15.65 % compared with control and nitroglycerine. Figure 1A

**Figure 1.**
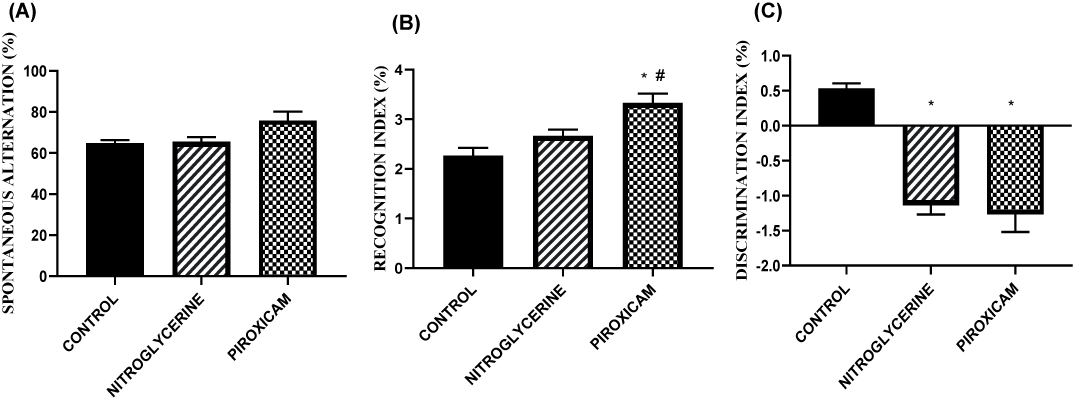
Effect of nitroglycerine and piroxicam on behavioral indices of learning and memory in female Wistar rats during di-oestrous. (A) Spontaneous alternation percentage evaluated via the Y-maze test. (B) Recognition Index (RI) percentages. (C) Discrimination Index (DI) percentages. Values are presented as Mean ± SEM (n = 5). Statistical significance is defined at p ≤ 0.05. *p < 0.05 compared with the control group; #p < 0.05 compared with the nitroglycerine group.

### Novel objects recognition test

#### Recognition index

Recognition Index in nitroglycerine-treated (2.67 ± 0.12 %) did not differ significantly when compared with control (2.27 ± 0.16 %) but decreased significantly when compared with piroxicam-treated (3.33 ± 0.19 %). The recognition index value in piroxicam-treated increased significantly compared with control value. Figure 1B

#### Discrimination index

Treatment with nitroglycerine (-1.138 ± 0.058 %) and piroxicam (-1.27 ± 0.011 %) significantly reduced the discrimination Index when compared with control (0.53 ± 0.03 %). Figure 1C

### Influences of nitroglycerine and piroxicam on neurotransmitters in di-oestrous rats

#### Glutamate level

Nitroglycerine treatment significantly reduced hippocampal glutamate level (42.50 ± 1.61µmol/g tissue) when compared with control (60.33 ± 5.81 µmol/g tissue) and piroxicam-treated (66.83 ± 9.82 µmol/g tissue). However, glutamate values in piroxicam-treated did not differ significantly when compared with control values. Figure 2A

**Figure 2.**
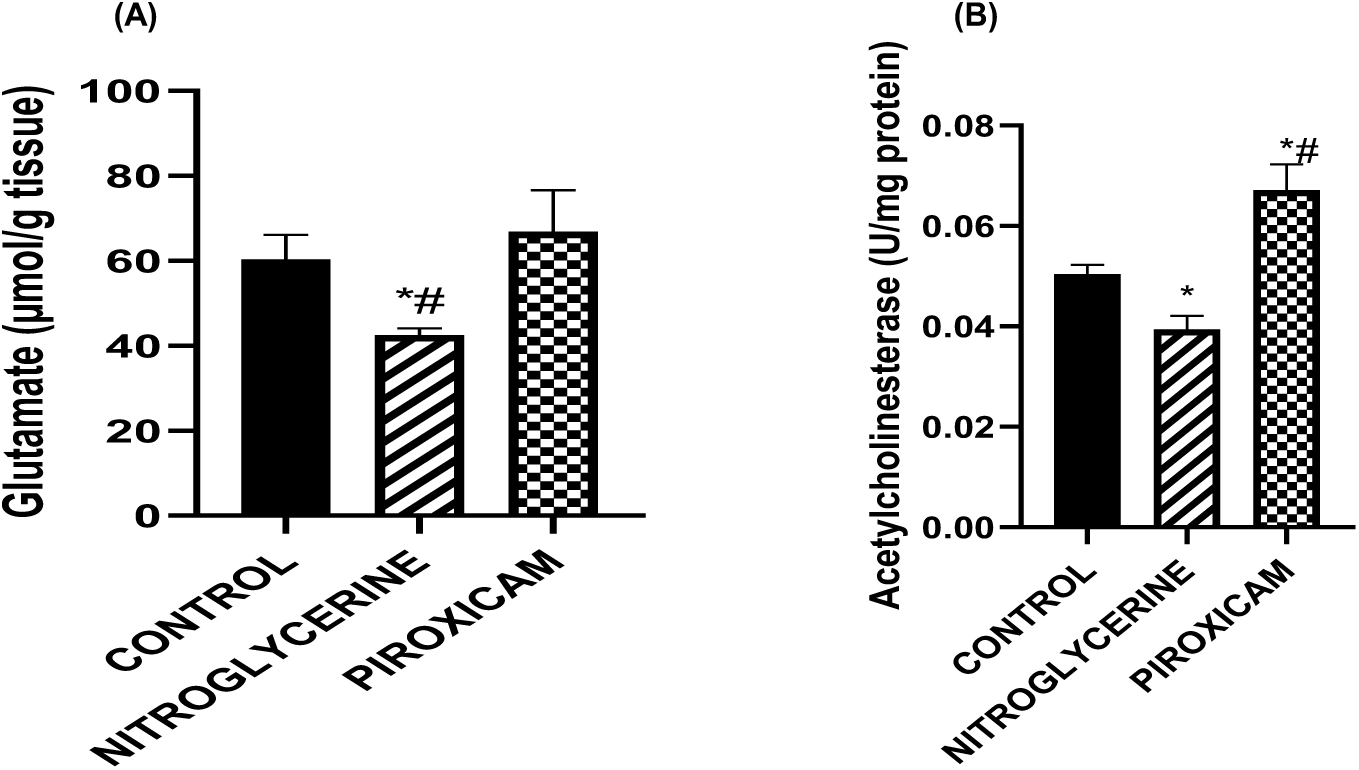
Neurochemical changes in hippocampal neurotransmission following nitroglycerine and piroxicam administration in female Wistar rats during di-oestrous. (A) Hippocampal glutamate levels. (B) Hippocampal acetylcholinesterase (AChE) levels. Values are presented as Mean ± SEM (n = 5). Statistical significance is defined at p ≤ 0.05. *p < 0.05 compared with the control group; #p < 0.05 compared with the Nitroglycerine group.

#### Acetylcholinesterase activity

Nitroglycerine treatment caused significant reduction in the activity of hippocampal acetylcholinesterase (AchE) (0.039 ± 0.003 U/mg protein) when compared with control. However, AchE activity was significantly increased following piroxicam treatment (0.067 ± 0.005 U/mg protein) when compared with control (0.050 ± 0.002 U/mg protein). Compared with piroxicam, nitroglycerine significantly reduced AchE activity. Figure 2B

### Influences of nitroglycerine and piroxicam on oxidative stress and inflammation in di-oestrous rats

#### Malondialdehyde level of female wistar rats during di-oestrous

Nitroglycerine treatment did not cause any significant difference in Malondialdehyde level (2.111 ± 0.1314 nmol/mg protein) when compared with control (2.235 ± 0.07676 nmol/mg protein). While piroxicam treatment (2.705 ± 0.04670 nmol/mg protein) significantly increased hippocampal malondialdehyde level compared with control. Also, administration of piroxicam significantly increased malondialdehyde level compared with nitroglycerine treatment. Figure 3A

**Figure 3.**
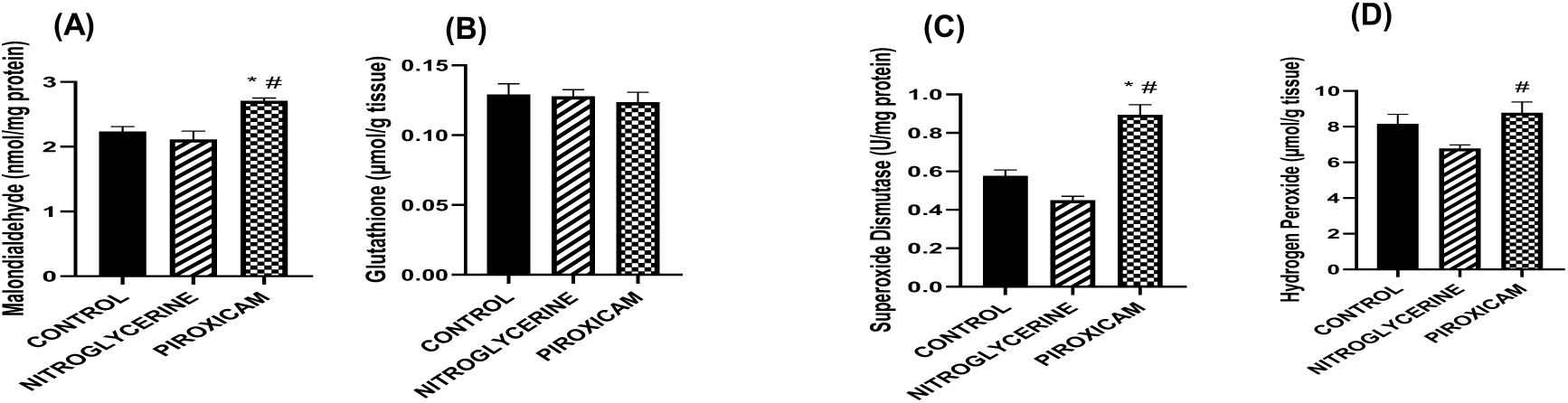
Hippocampal oxidative and antioxidative stress biomarker levels during di-oestrous in female Wistar rats treated with nitroglycerine and piroxicam. (A) Hippocampal malondialdehyde (MDA) levels. (B) Hippocampal glutathione (GSH) levels. (C) Hippocampal superoxide dismutase (SOD) levels. (D) Hippocampal hydrogen peroxide (H2O2) levels. Values are presented as Mean ± SEM (n = 5). Statistical significance is defined at p ≤ 0.05. *p < 0.05 compared with the control group; #p < 0.05 compared with the nitroglycerine group.

#### Glutathione level

Treatment with nitroglycerine and piroxicam (0.1278 ± 0.0048 and 0.1236 ± 0.0071 µmol/g tissue, respectively) did not cause significant difference in hippocampal glutathione level when compared with control (0.1290 ± 0.0078 µmol/g tissue). Figure 3B

#### Superoxide dismutase activity

Treatment with nitroglycerine did not cause any significant difference in hippocampal superoxide dismutase activity (0.449 ± 0.021 U/mg protein) when compared with control (0.576 ± 0.031 U/mg protein). While piroxicam treatment (0.895 ± 0.052 U/mg protein) significantly increased hippocampal superoxide dismutase level compared with control. Also, administration of piroxicam significantly increased superoxide dismutase level compared with nitroglycerine treatment. Figure 3C

#### Hydrogen peroxide level

Nitroglycerine and piroxicam treatment (6.778 ± 0.201and 8.783 ± 0.601 µmol/g tissue, respectively) did not cause any significant difference in hippocampal hydrogen peroxide level when compared with control (8.17 ± 0.53 µmol/g tissue). However, Piroxicam treatment significantly increased hydrogen peroxide level compared with nitroglycerine. Figure 3D

#### Myeloperoxidase activity

Nitroglycerine and piroxicam treatment significantly increased myeloperoxidase activity (0.1066 ± 0.0021 U/mg tissue and 0.089 ± 0.0039 U/mg tissue respectively) when compared with control (0.0250 ± 0.0026 U/mg tissue). Nitroglycerine treatment also increased myeloperoxidase activity significantly by 326.4 % and 19.7 % compared with control and piroxicam treated. Figure 4A

**Figure 4.**
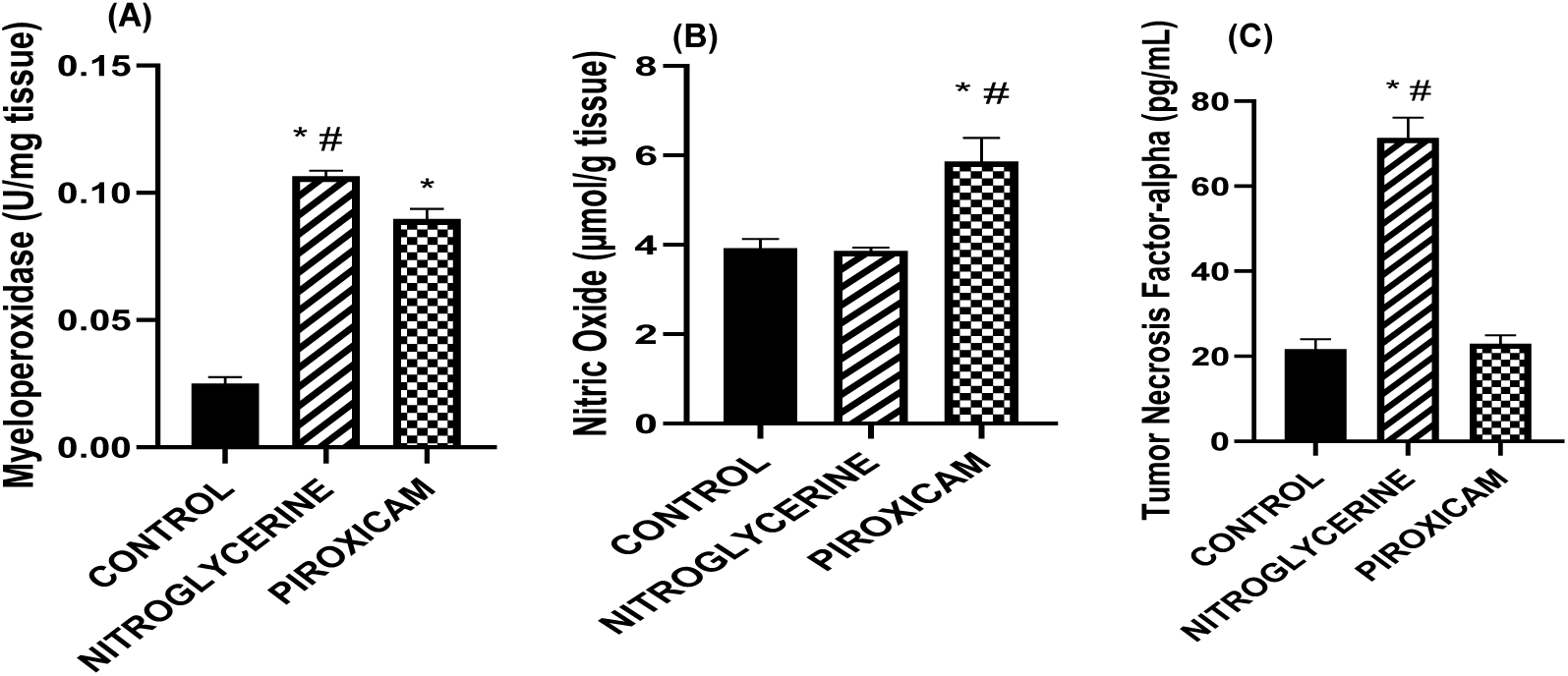
Hippocampal neuroinflammatory and nitrosative biomarker profiles during di-oestrous following treatment. (A) Hippocampal myeloperoxidase (MPO) levels. (B) Hippocampal nitric oxide (NO) levels. (C) Hippocampal tumor necrosis factor-alpha (TNF-α) levels. Values are presented as Mean ± SEM (n = 5). Statistical significance is defined at p ≤ 0.05. *p < 0.05 compared with the control group; #p < 0.05 compared with the comparative treatment group (piroxicam or nitroglycerine respectively).

#### Nitric oxide level

Treatment with nitroglycerine did not cause any significant change in nitric oxide level (3.86 ± 0.077 µmol/g tissue) when compared with control (3.926 ± 0.202 µmol/g tissue). While piroxicam treatment caused a significant increase in nitric oxide level (5.86 ± 0.53 µmol/g tissue) compared with control. Administration of piroxicam significantly increased nitric oxide level compared with nitroglycerine treatment. Figure 4B

#### Tumour necrosis factor-alpha level

Nitroglycerine treatment caused a significant increase in TNF-α level (71.38 ± 4.81 pg/mL) compared with control and treatment with piroxicam (21.70 ± 2.34 and 23.00 ± 1.95 respectively pg/mL). Piroxicam treatment did not cause any significant difference in TNF-α level compared with control. Figure 4C

### Effect of nitroglycerine and piroxicam on hippocampal histoarchitecture in di-oestrous rats

#### Control group

Histological examination of H&E-stained hippocampal sections from the control group revealed normal cytoarchitecture, with well-preserved pyramidal and granule cell layers. The hippocampal architecture appeared intact, with no evidence of neuronal loss, structural disruption, or other histopathological abnormalities. Figure 5A

**Figure 5.**
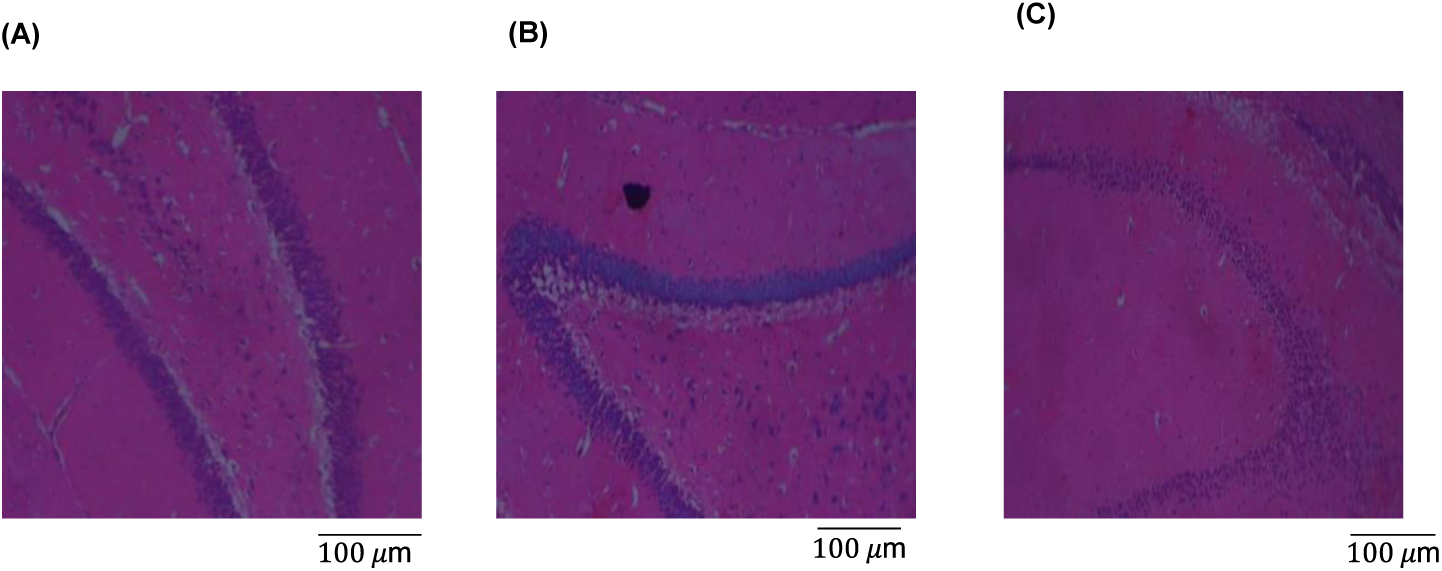
Photomicrographs of hematoxylin and eosin (H&E) stained hippocampal sections (magnification ×100) in female Wistar rats during di-oestrous. (A) Control group, (B) Nitroglycerine-treated group, and (C) Piroxicam-treated group, all exhibiting normal histological and cellular architecture. Scale bar = 100 μm.

#### Nitroglycerine-treated group

H&E-stained sections from the nitroglycerine-treated group demonstrated preservation of the normal hippocampal architecture. The pyramidal and granule cell layers remained continuous and well organized, with no evident neuronal degeneration, tissue distortion, or significant histopathological alterations when compared with the control group. Figure 5B

#### Piroxicam-treated group

Histological evaluation of the piroxicam-treated group showed intact hippocampal cytoarchitecture, characterised by preserved pyramidal and granule cell layers. No appreciable neuronal loss, structural abnormalities, or other overt histopathological changes were observed relative to the control animals. Figure 5c

## Discussion

This study investigated the effects of nitroglycerine and piroxicam on hippocampal-dependent cognitive performance and neurochemical indices in di-oestrous female Wistar rats, a hormonal phase associated with increased inflammatory sensitivity. The findings demonstrate that both agents exert distinct central effects on memory-related behaviour and hippocampal biochemical markers, despite preserving basic spatial working memory. The Y-maze and NORT assessed spatial working and recognition memory, respectively, with spontaneous alternation, recognition index, and discrimination index serving as behavioural measures of hippocampal function [19–22]. Reduced index score indicates hippocampal dysfunction [23].

In this study, while spontaneous alternation did not differ in both piroxicam-treated and nitroglycerine groups relative to control, indicating preservation of short-term spatial working memory and gross hippocampal integrity that suggests neither treatment induced overt cognitive toxicity or widespread neuronal dysfunction, the recognition index for piroxicam treated was significantly improved but this did not translate to improved memory precision, as the discrimation index reduced following both piroxicam and nitroglycerine treatment. This could be indicative of partial hippocampal dysfunction, as well as reduced learning and memory consolidation. Consolidation, which refers to the process of converting temporary memories into long-lasting ones [24], may have been impaired. This dissociation supports the concept that object recognition and discrimination depend on partially distinct neural processes, with discrimination reflecting finer hippocampal–perirhinal integration and higher demands on synaptic plasticity. Thus, the observed behavioural pattern suggests qualitative alterations in memory processing rather than complete impairment of memory formation. It has been established that the role of hippocampus in memory function is often regulated by prostanoids, such as prostaglandins. Prostaglandins play a regulatory role in several forms of neural plasticity, including long-term potentiation, a cellular model for certain forms of learning and memory [25]. Piroxicam, as an over-the-counter drug used to relieve dysmenorrhea, by inhibiting COX activity that is upregulated during menstruation, can impact memory potentiation which was observed with the reduced discriminative index in this study. Piroxicam and nitroglycerine treatment may have also reduced hippocampal COX activity and/or prostanglandin-receptor binding to alter consolidation of learning and memory. Piroxicam may have enhanced recognition memory through its anti-inflammatory and antioxidant effects [26] but not the ability to differentiate between objects, which indicates impaired novelty-based decision-making. The reduced discrimaination index with non-differential change in spontatenous alternation and recognition index following nitroglycerine treatment indicates that nitroglycerine may not have a direct impact on spatial memory and recognition memory but adversely affected precision with decision making alongside piroxicam.

Inflammation in the central nervous system has been linked to various neurodegenerative diseases, with evidence showing that reducing inflammation can enhance cognitive function [27]. Along with other factors such as peripheral inflammation, impaired cerebral blood flow, oxidative stress etc., inflammation may affect memory at the different phases [28]. Neutrophils, the most abundant peripheral immune cells, respond to infection by adhering to blood vessel walls and migrating to site of infection. Increased penetration of the brain by neutrophils arises from increased disruption of blood-brain barrier consequent of increased peripheral inflammatory response. Enhanced neutrophil extravasation into the central nervous system has been reported to contribute to disorders of the CNS [29]. Myeloperoxidase, the most abundant protein in neutrophils, mediates neutrophils oxidative response to promote tissue oxidative damage and inflammation [30]. Increasing activity of hippocampal myeloperoxidase with piroxicam and nitroglycerine treatments is indicative of neuroinflammation promotion which may have impacted consolidation of memory and learning resulting in reduced discriminative index. Piroxicam has been reported to possess anti-inflammatory effects [31], but the observed reduced discriminative index did not translate to this.

The tumour necrosis factor-alpha (TNF-α) level was significantly elevated in the nitroglycerine-treated group compared with both control and piroxicam-treated groups. TNF-α is a pro-inflammatory cytokine associated with neuroinflammation and cognitive decline [32]. The increase in TNF-α in nitroglycerine-treated rats aligns with studies suggesting that nitric oxide donors can induce neuroinflammatory responses [33]. Nitric oxide signaling has been implicated in the activation of inflammatory transcription pathways, including NF-κB, which regulates pro-inflammatory cytokine expression [34]. The selective elevation of TNF-α without accompanying oxidative damage suggests a predominantly cytokine-driven inflammatory response rather than a reactive oxygen species–dominated process.

The results showed no significant change in hippocampal nitric oxide levels in the nitroglycerine-treated group compared to control, whereas a significant increase was observed in the piroxicam-treated group. Nitric oxide is a signaling molecule involved in neurotransmission and inflammatory regulation [35]. While basal nitric oxide levels may exert modulatory roles, excessive nitric oxide production can promote nitrosative stress through the formation of reactive nitrogen species such as peroxynitrite [36]. The elevated nitric oxide levels observed following piroxicam treatment, together with increased malondialdehyde and myeloperoxidase activity, suggest enhanced nitrosative and oxidative stress rather than an anti-inflammatory effect. This oxidative–inflammatory environment may contribute to the increased MPO levels observed in the piroxicam-treated group.

Oxidant/antioxidant balance in this study, can be reflected by the level of malondialdehyde, a marker of lipid peroxidation and oxidative stress, and activity of antioxidant enzymes [37]. The imbalance has been associated with various neurodegenerative diseases and cognitive impairment. Prolonged oxidative response consequently leads to inflammatory response that could become sustained and detrimental [30]. Findings from this study showed increased malondialdehyde level and superoxide dismutase (SOD) activity with piroxicam treatment while nitroglycerine showed non-differential effect on malondialdehyde and antioxidant enzyme activity suggesting preservation of the antioxidant defence system. This activity of nitroglycerine in preserving the antioxidant defence system could also be linked with its reported antioxidant potentials [38]. The increase in SOD activity with piroxicam treatment suggests a possible compensatory mechanism for the increased level of malondialdehye to prevent oxidative damage [39]. This could also be advanced for the non-differential increase in peroxide level. Hydrogen peroxide, a necessary signalling molecule at normal levels but a potential contributor to neurodegenerative diseases when excessively accumulated [40]. The absence of significant changes in glutathione and hydrogen peroxide levels across groups suggests that oxidative stress was moderate and localized rather than widespread.

Glutamate, a key excitatory neurotransmitter [41], plays a critical role in synaptic plasticity, learning and memory formation [42]. Disruptions in glutamatergic signaling can adversely affect these cognitive functions [43]. The significant reduction in glutamate levels in the nitroglycerine-treated group suggests that nitroglycerine may impair glutamatergic neurotransmission, potentially contributing to cognitive deficits and the observed discriminative index in this study. Impaired glutamatergic neurotransmission, which involves disruptions in the signalling of glutamate, the brain’s primary excitatory neurotransmitter has been implicated in various neurological and psychiatric disorders [44]. Although N-methly-D-aspartate (NMDA) receptor activity was not directly assessed, reduced glutamate availability is consistent with diminished glutamatergic drive, which may preserve baseline cognitive performance while impairing memory precision. In addition, acetylcholinesterase (AChE) activity was also reduced by nitroglycerine treatment. AChE is an enzyme that breaks down acetylcholine [45], a neurotransmitter involved in learning and memory [46]. The reduction in AChE activity in the nitroglycerine group may lead to an accumulation of acetylcholine, potentially enhancing cholinergic transmission [47]. However, this effect did not translate into improved cognitive performance in the Y-Maze or NORT, with recent researches suggesting that Elevated hippocampal acetylcholine promotes maladaptive stress-related behaviours [48]. In contrast, piroxicam did not alter glutamate levels but increased AChE activity suggesting that its antioxidant and anti-inflammatory properties may have helped to maintain normal glutamate homeostasis but increased acetylcholine breakdown. The increase in AChE activity and unaltered glutamate level could be linked to the observed improvements in the recognition memory [49]. However elevated AChE activity has been implicated in various neurodegenerative diseases including Alzheimer’s disease [50]. Given the established role of acetylcholine in attention, encoding, and hippocampal-dependent memory accuracy, reduced cholinergic signaling may contribute to the impaired discrimination index observed in the piroxicam-treated group, despite improved recognition performance. H&E staining revealed preserved hippocampal architecture in all groups, indicating that the observed behavioural and biochemical changes occurred in the absence of overt structural damage.

## Conclusions

In conclusion, this study demonstrates that nitroglycerine and piroxicam exert distinct central effects on hippocampal neurochemistry and memory processing in our di-oestrous–based inflammatory model. Nitroglycerine predominantly induces neuroinflammatory modulation with preserved redox balance, while piroxicam promotes oxidative and nitrosative stress–linked hippocampal alterations despite partial cognitive benefits. These findings emphasize the importance of considering central nervous system consequences when evaluating analgesic interventions in dysmenorrhea and related inflammatory conditions, and they contribute to a growing body of evidence that pain management strategies may influence cognitive function through differential neurochemical pathways.

